# Delirium, frailty and mortality: interactions in a prospective study of hospitalized older people

**DOI:** 10.1101/207621

**Authors:** Melanie Dani, Lucy H Owen, Thomas A Jackson, Kenneth Rockwood, Elizabeth L Sampson, Daniel Davis

## Abstract

**Background:** It is unknown if the association between delirium and mortality is consistent for individuals across the whole range of health states. A bimodal relationship has been proposed, where delirium is particularly adverse for those with underlying frailty, but may have a smaller effect (perhaps even protective) if it is an early indicator of acute illness in fitter people. We investigated the impact of delirium on mortality in a cohort simultaneously evaluated for frailty.

**Methods:** We undertook an exploratory analysis of a cohort of consecutive acute medical admissions aged ≥70. Delirium on admission was ascertained by psychiatrists. A Frailty Index (FI) was derived according to a standard approach. Deaths were notified from linked national mortality statistics. Cox regression was used to estimate associations between delirium, frailty and their interactions on mortality.

**Results:** The sample consisted of 710 individuals. Both delirium and frailty were independently associated with increased mortality rates (delirium: HR 2.4, 95%CI 1.8-3.3, p<0.01; frailty (per SD): HR 3.5, 95%CI 1.2-9.9, p=0.02). Estimating the effect of delirium in tertiles of FI, mortality was greatest in the lowest tertile: tertile 1 HR 3.4 (95%CI 2.1-5.6); tertile 2 HR 2.7 (95%CI 1.5-4.6); tertile 3 HR 1.9 (95% CI 1.2-3.0).

**Conclusion:** While delirium and frailty contribute to mortality, the overall impact of delirium on admission appears to be greater at lower levels of frailty. In contrast to the hypothesis that there is a bimodal distribution for mortality, delirium appears to be particularly adverse when precipitated in fitter individuals.

## Introduction

Delirium, characterized by a fluctuating disturbance in arousal, attention and cognition secondary to an acute medical condition, is common, affecting 18-35% of general medical inpatients, 8-17% of older patients attending emergency departments and 51% of patients in post-acute care.(1-3) It is associated with increased length of stay, institutionalization, and progression of dementia.(4-7)

Delirium is widely understood to be associated with mortality, with an overall HR=1.9 consistent across a number of studies identified in a systematic review.(4) This meta-analysis included observational studies adjusting for chronic comorbidity or acute illness, though none accounted for both. Therefore, an important unanswered question is whether the association remains independent of acute *and* chronic health factors that might otherwise drive mortality.(8) Indeed, a more nuanced understanding of delirium and mortality is relevant given the proposal that the relationship may be bimodal.(9) That is, while delirium may have catastrophic outcomes in some patients, for others, it may be an early indicator of acute illness leading to earlier recognition and treatment, perhaps even being protective.

Accounting for underlying frailty may provide an insight into the relationship between delirium and mortality because frailty itself is so closely related to risk of both mortality and delirium. One view of frailty describes the gradual accumulation of deficits as individuals age, which results in loss of physiological reserve (physical, mental and functional) and increased vulnerability to insults.(10) Taking baseline functional status and chronic comorbidity into account can resolve the issue of ‘unmeasured heterogeneity’ - the factors which increase risk despite the same level of acute illness.(11)

Our aim was to investigate the effect of delirium and frailty on mortality in a large cohort of acutely unwell adults, setting out to answer the following questions: (1) is delirium associated with mortality, even after adjusting for underlying frailty? (2a) is there an interaction between delirium and frailty? (2b) is the relationship with frailty and mortality linear across the range of frailty states and does this change according to the presence of delirium?

## Methods

We undertook an exploratory analysis of a cohort prospectively ascertaining outcomes from acutely hospitalized elders. Participants were recruited as previously described.(12) Briefly, all patients aged ≥70 years consecutively admitted to the acute medical unit between June 2007 and December 2007 were screened for inclusion. Exclusion criteria were 1) admission length of less than 48 hours and 2) insufficient English to be assessed for cognitive and mental status. The Royal Free Hospitals NHS Trust Ethics Committee gave ethical approval (06/Q0501/31).

## Outcomes

The study was notified of all deaths through linkage with the UK Office for National Statistics for up to 3 years following the index admission.

## Exposures

### Delirium

All participants were evaluated on admission by trained psychiatrists. Formal cognitive testing included the Mini-Mental State Examination(13) and delirium was defined using the Confusion Assessment Method (CAM) algorithm.(14) Key symptoms such as inattention were identified through the “serial 7s” or “W-O-R-L-D backwards” tasks. Other items such as acute onset, disorganized thinking and altered level of consciousness, degree of fluctuation of these symptoms, were ascertained through clinical assessment which included information from ward staff and the medical chart.

### Frailty

A 31-item FI was constructed according to a standardized procedure, (15) where the following variables were included: comorbidity (13 variables), examination findings (5 variables), laboratory findings (9 variables) and functional status (4 variables) (Table 1). These variables were selected to encompass the full range of acute and chronic health factors that could account for any observed association with mortality. All items were given a binary score (0 = no deficit, 1 = deficit present). For each participant, an FI score was calculated by dividing the number of deficits present by the denominator of 31 maximum deficits, resulting in a score between 0 and 1. For example, for an individual with 10 deficits present out of 31, their FI score would be 10/31=0.32. Across several iterations of FI in several hundred datasets, the usual upper limit of frailty observed asymptotically approaches 0.70. In our dataset, data were not missing for more than 6% in each variable.

**Figure 1.**
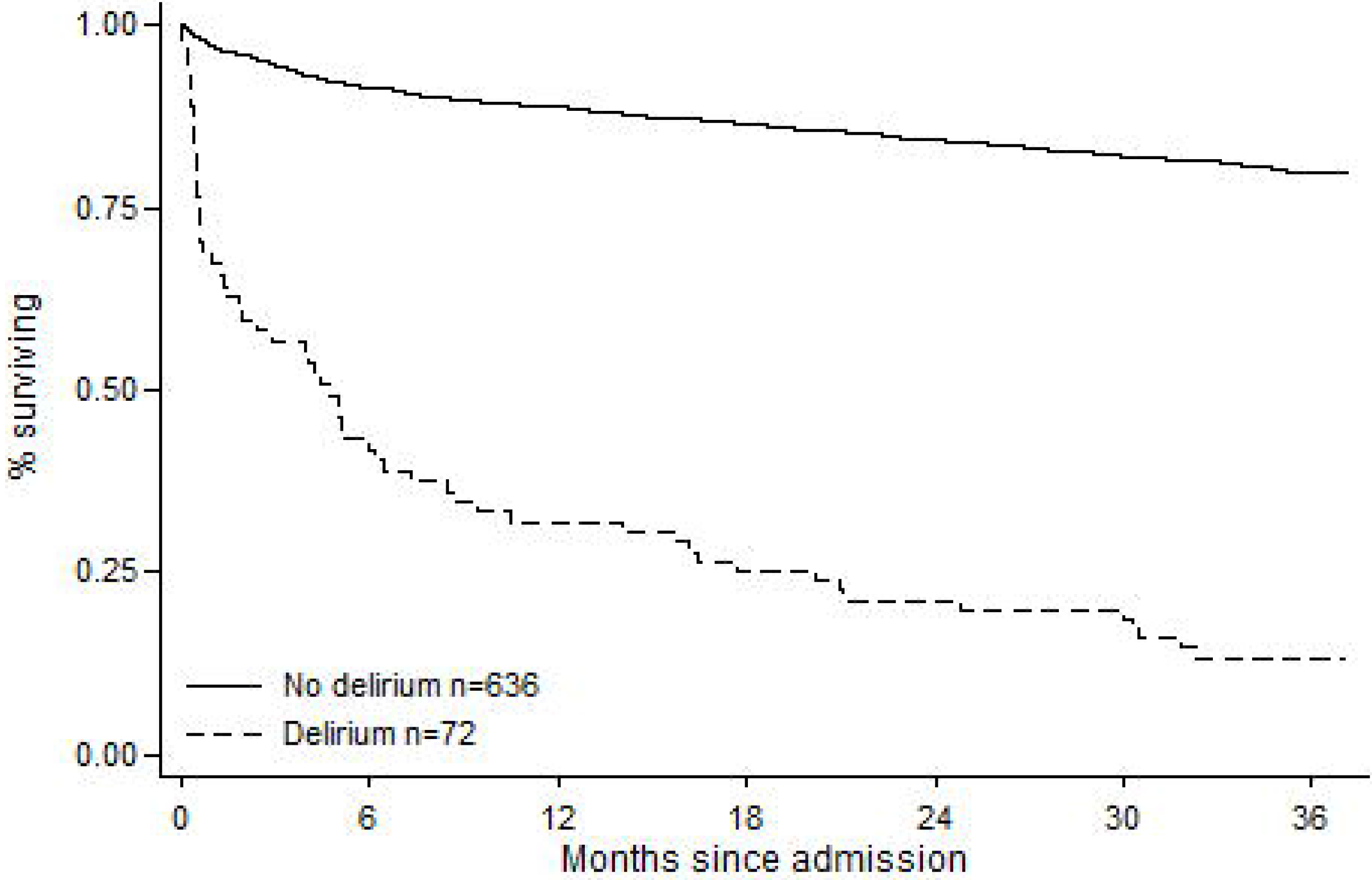
Kaplan-Meier curves showing survival of cohort, by delirium status, adjusted by age, sex and FI score.

### Statistical analysis

Proportional hazards for mortality were assessed in a series of Kaplan-Meier plots and Cox regression models, where outcome was date of death. Post-estimation procedures included Schoenfeld residuals for checking assumptions of proportionality. Multiplicative interactions between delirium and FI score were assessed in order to estimate the association of delirium and mortality with respect to underlying frailty, using α=0.1 as a threshold for type 1 errors. This approach has been justified by quantifying the gains in power using a less stringent α for samples of this size where interactions between dichotomous (delirium status) and continuous (frailty index) variables are being considered.(16) Linearity of any associations with mortality across the distribution of frailty, restricted cubic splines were fitted (four knots), plotting log-hazard ratio for mortality against FI score, stratified by delirium status. Stata version 14.1 was used for all statistical procedures.

## Results

The sample contained 710 individuals, with a mean age of 83.1 years (standard deviation 7.41), and 59% were female (Supplementary Table 1). A diagnosis of dementia was present in 42%. At the end of the three-year follow-up, 340 individuals remained alive and 370 had died (median follow-up 5 months, IQR 1 to 17 months). The prevalence of delirium on admission was 10.3% (n=73). The prevalence of deficits on index items ranged from 6% (metastatic disease) to 73% (CRP>5) (Table 1). The mean FI score was 0.23 (SD 0.096, upper limit 0.55), with a broadly normal distribution (Supplementary Figure 1).

**Table 1.**
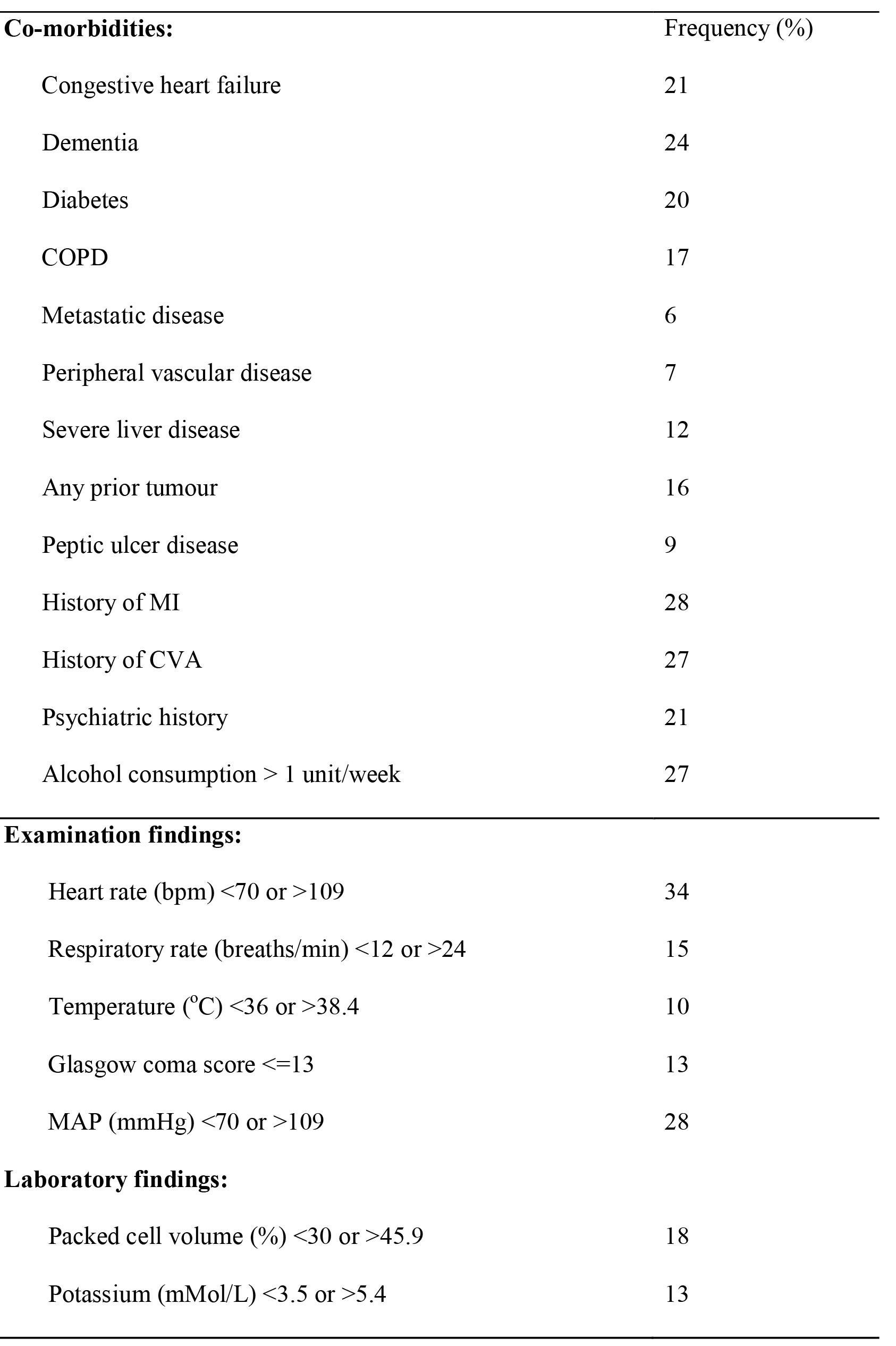
Frailty index variables

| **Co-morbidities:** | Frequency (%) |
| --- | --- |
| Congestive heart failure | 21 |
| Dementia | 24 |
| Diabetes | 20 |
| COPD | 17 |
| Metastatic disease | 6 |
| Peripheral vascular disease | 7 |
| Severe liver disease | 12 |
| Any prior tumour | 16 |
| Peptic ulcer disease | 9 |
| History of MI | 28 |
| History of CVA | 27 |
| Psychiatric history | 21 |
| Alcohol consumption > 1 unit/week | 27 |
| **Examination findings:** |  |
| Heart rate (bpm) <70 or >109 | 34 |
| Respiratory rate (breaths/min) <12 or >24 | 15 |
| Temperature (°C) <36 or >38.4 | 10 |
| Glasgow coma score ≤13 | 13 |
| MAP (mmHg) <70 or >109 | 28 |
| **Laboratory findings:** |  |
| Packed cell volume (%) <30 or >45.9 | 18 |
| Potassium (mMol/L) <3.5 or >5.4 | 13 |

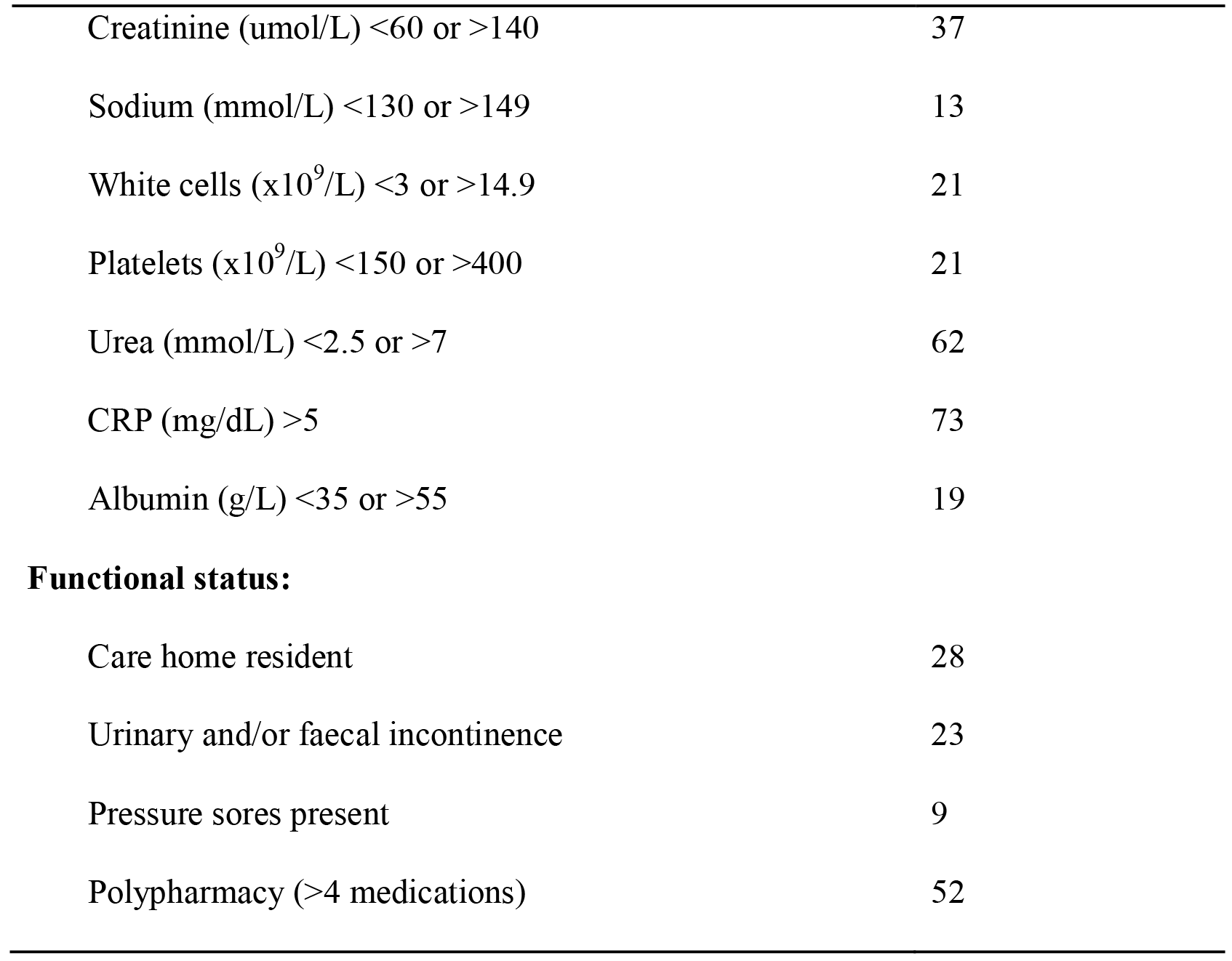

|  |  |
| --- | --- |
| Creatinine (umol/L) <60 or >140 | 37 |
| Sodium (mmol/L) <130 or >149 | 13 |
| White cells ($\times 10^9$/L) <3 or >14.9 | 21 |
| Platelets ($\times 10^9$/L) <150 or >400 | 21 |
| Urea (mmol/L) <2.5 or >7 | 62 |
| CRP (mg/dL) >5 | 73 |
| Albumin (g/L) <35 or >55 | 19 |
| **Functional status:** |  |
| Care home resident | 28 |
| Urinary and/or faecal incontinence | 23 |
| Pressure sores present | 9 |
| Polypharmacy (>4 medications) | 52 |

### Delirium, frailty and mortality

Delirium was strongly associated with mortality, with 59 (81%) dying within the next three years, compared with 311 (49%) without delirium. Tertiles of FI score (first 0-0.19; second 0.20- 0.26; third (0.27- 0.55) also showed increasing associations with mortality (first 48%; second 51%; third 60%). Both delirium and frailty were crudely associated with increased hazard for death (delirium: 2.4, 95% CI 1.8- 3.2; FI (per SD); HR 5.9 (95% CI 2.1-16)). This remained the case with a multivariable model including both terms adjusted by age and sex (delirium: HR 2.4, 95% CI 1.8-3.3, p<0.01; frailty (per SD): HR 3.5, 95% CI 1.2-9.9, p=0.02) (Table 2). Kaplan-Meier survival curves (adjusted by age and sex), according to delirium status demonstrated worse survival for those with delirium (Figure 1).

**Table 2.** Survival analysis showing the associations between delirium, frailty and mortality.

|  | Univariable |  |  |  | Multivariable |  |  |  |
| --- | --- | --- | --- | --- | --- | --- | --- | --- |
|  | Hazard<br>ratio | 95% CI |  | p | Hazard<br>ratio | 95% CI |  | p |
| Delirium | 2.44 | 1.85 | 3.23 | <0.01 | 2.37 | 1.78 | 3.15 | <0.01 |
| Frailty<br>Index | 5.90 | 2.14 | 16.2 | <0.01 | 3.37 | 1.18 | 9.60 | 0.02 |
| N=708, all models adjusted by age and sex. Mean FI score = 0.23 (SD 0.09) |  |  |  |  |  |  |  |  |

### Interactions between delirium and frailty

We estimated the effect of delirium on mortality according to each tertile of FI. There was an inverse gradient of association, where stronger associations were observed in the fittest group (tertile 1 HR 3.4 (95%CI 2.1-5.6); tertile 2 HR 2.7 (95%CI 1.5 -4.6); tertile 3 HR 1.9 (95% CI 1.2-3.0)). The delirium-frailty interaction was statistically significant if α=0.1 (p=0.07).

### Linearity of mortality gradients in relation to delirium status

Restricted cubic splines of the log-HR for mortality against FI score were linear across the four knots, giving no indication that mortality is driven by a particular portion of the frailty continuum (Figure 2). When additionally accounting for delirium status, the splines remained linear, though crossed over to suggest delirium may have greater associations with mortality at lower degrees of frailty and lower associations at higher degrees of frailty (p=0.07) (Figure 2).

**Figure 2.**
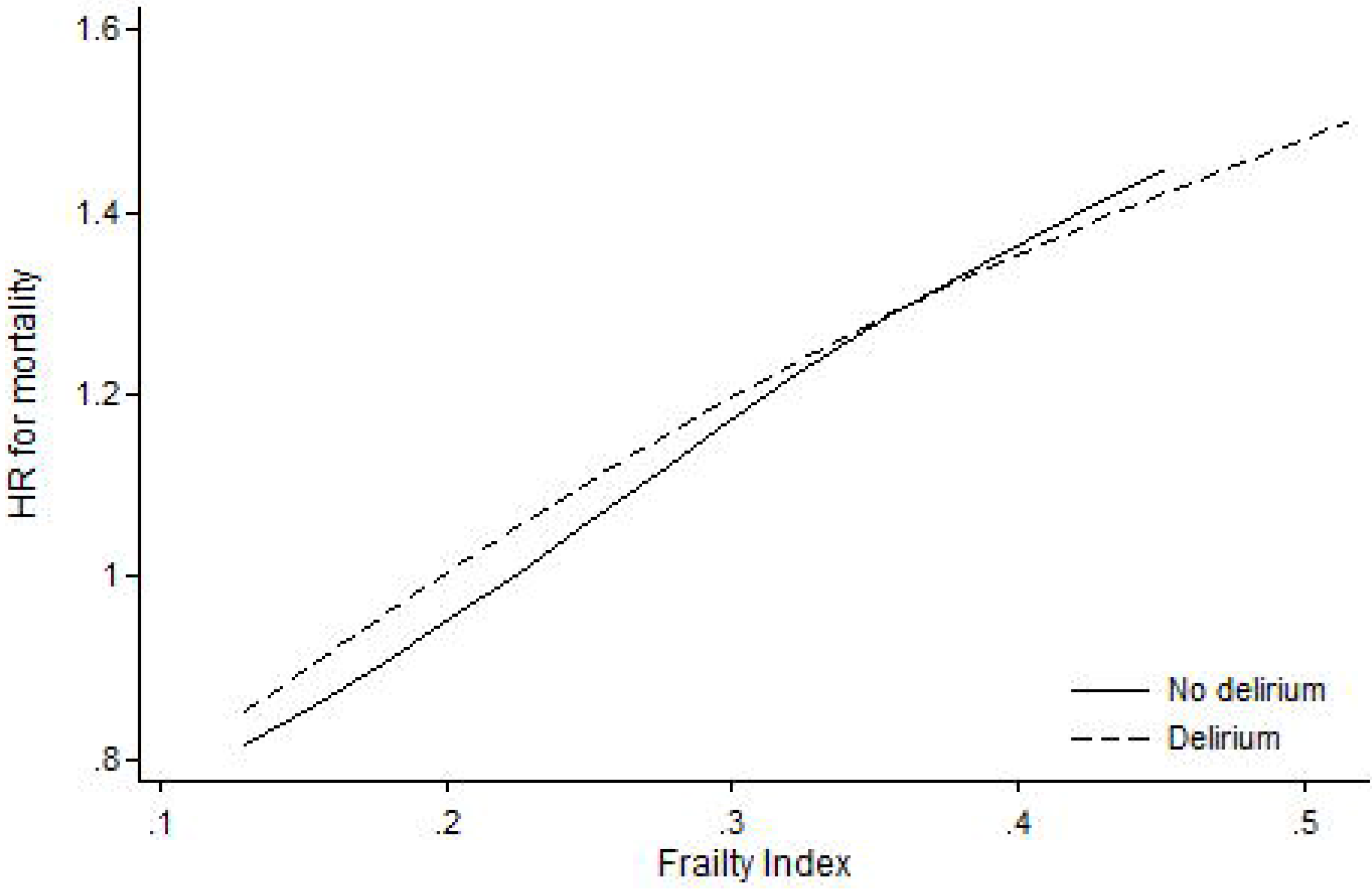
Relationship between frailty and mortality, by delirium status Restricted cubic splines modelling relationship between frailty and mortality. The linearity suggests a continuous relatinoship between frailty and mortality. When stratified by delirium, the lines intersect suggesting slightly greater effects with delirium at lower levels of mortality (p=0.07)

## Discussion

We found that on admission, both delirium and frailty were independently associated with increased risk of death, beyond that expected from all the acute and chronic health factors measured in this study. Moreover, delirium and frailty appeared to interact in a way that while delirium increases the risk of death at all levels of frailty, the relative impact of this association was greatest in fitter patients. The relationship between frailty and mortality was linear, suggesting mortality is not being driven by a subset of individuals. Delirium status had a small influence on these relationships. Taken together, our results suggest that delirium itself independently confers a mortality risk, and this risk applies whatever he underlying degree of frailty. Our data illustrate some of the advantages of using a frailty index in studying cognitive disorders by quantifying a multidimensional assessment, thereby allowing other ‘non-neurological factors’ that nevertheless contribute to patient outcomes (for example, nutritional status, polypharmacy, electrolyte abnormalities) to be considered.(17)

Our findings should be treated with caution. Data were collected from a single site, albeit a large London secondary care hospital with unselected medical admissions from a population of 1.2 million people and high generalizability. Secondly, despite a standardized protocol and training procedure, more than one rater performed the psychiatric evaluation, possibly introducing inter-rater variability. Thirdly, patients were assessed for delirium on admission only (defined as within 72 hours of admission). Any patients developing delirium after this point would not be included in this analysis, possibly underestimating the effects of delirium. Strengths included large sample size and validated assessments undertaken by specialist diagnosticians. Moreover, a wide variety of variables were available to construct a broad FI. The specific inclusion of physiological and laboratory items may generally relate to acute illness, it is becoming clear that use of such measures are nonetheless informative for use in a frailty index.(18) Though functional items were under-represented, for the requirements of this specific analysis, the combination of acute and chronic health factors made for a particularly robust FI measure.

To understand why the relative risk is higher in fit patients compared with frail patients, the insult causing delirium in fit individuals needs to be large, in order to overcome their physiological and cognitive reserve. Another reason for delirium in fitter individuals may be a distinct neurological precipitant which could drive the poorer prognosis in these patients. Both frailty and delirium are associated with vulnerability and lack of physiological and cognitive reserve to insults, though few studies have been able to address this directly.(19) One study found that individuals without a prior diagnosis of dementia (a marker of frailty and reserve) had a worse prognosis from delirium than those with a diagnosis, supporting our finding that fitter patients may incur a worse prognosis.(20) A separate finding that mortality after delirium appears to arise independently of illness severity or frailty.(21) We report a similar distribution of Frailty Index scores, though our study was large enough to explore the interactions presented here.

The mechanisms by which delirium independently increases risk of death (after adjusting for illness severity) remain unclear. Delirium complicates and impairs recovery – patients with delirium are more likely to receive psychotropic drugs, more likely to fall and less likely to mobilize effectively during and after their illness, all of which may have an adverse impact on survival.(22) In addition, they may be less likely to maintain adequate hydration and malnutrition, and be less compliant with medication. Of particular concern might be hypoactive delirium, where reduced arousal might lead to aspiration pneumonia. Overall, such findings serve to emphasize the emergency nature of delirium.

To conclude, in this exploratory analysis of a large cohort of consecutive admissions to an acute adult medical unit, we found that both delirium and frailty independently increase the risk of death. In addition, the risk of death is higher in delirious patients at all levels of frailty. The most striking finding using this approach is that the risk of death from delirium is highest in the fittest patients. This highlights the crucial importance of preventing, detecting and treating delirium in any patient, and recognizing it as a serious condition with prognostic significance.

## Captions

Supplementary Table 1: Cohort characteristics

Supplementary Figure 1. Distribution of a frailty index incorporating acute and chronic health factors.

## References

1. Siddiqi N, House AO, Holmes JD. Occurrence and outcome of delirium in medical in-patients: a systematic literature review. Age and Ageing. 2006;35:350–364.

2. Kiely DK, Bergmann MA, Jones RN, Murphy KM, Orav EJ, Marcantonio ER. Characteristics associated with delirium persistence among newly admitted post-acute facility patients. J Gerontol A Biol Sci Med Sci. 2004;59:344–349.

3. Jackson TA, Gladman JRF, Harwood RH, MacLullich AMJ, Sampson EL, Sheehan B, et al. Challenges and opportunities in understanding dementia and delirium in the acute hospital. PLoS medicine. 2017;14:el002247.

4. Witlox J, Eurelings LSM, DeJonghe JFM, Kalisvaart P, Eikelenboom P, VanGool WA. Delirium in Elderly Patients and the Risk of Postdischarge Mortality, Instituationalisation and Dementia. Journal of the American Medical Association. 2010;304:443–451.

5. Gross AL, Jones RN, Habtemariam DA, Fong TG, Tommet D, Quach L. et al. Delirium and long-term cognitive trajectory among persons with dementia. Archives of internal medicine. 2012;172:1324–1331.

6. Davis DH, Muniz-Terrera G, Keage HA, Stephan BC, Fleming J, Ince PG,et al. Association of Delirium With Cognitive Decline in Late Life: A Neuropathologic Study of 3 Population-Based Cohort Studies. JAMA psychiatry. 2017;74:244–251.

7. Davis DH, Muniz Terrera G, Keage H, Rahkonen T, Oinas M, Matthews FE,et al. Delirium is a strong risk factor for dementia in the oldest-old: a population-based cohort study. Brain : a journal of neurology. 2012;135:2809–2816.

8. Bellelli G, Frisoni GB, Turco R, Lucchi E, Magnifico F, Trabucchi M. Delirium superimposed on dementia predicts 12-month survival in elderly patients discharged from a postacute rehabilitation facility. J Gerontol A Biol Sci Med Sci. 2007;62:1306–1309.

9. Jackson TA, Wilson D, Richardson S, Lord JM. Predicting outcome in older hospital patients with delirium: a systematic literature review. Int J Geriatr Psychiatry. 2016;31:392–399.

10. Kelaiditi E, Andrieu S, Cantet C, Vellas B, Cesari M, Group ID. Frailty Index and Incident Mortality, Hospitalization, and Institutionalization in Alzheimer's Disease: Data From the ICTUS Study. J Gerontol A Biol Sci Med Sci. 2016;71:543–548.

11. Rockwood K, Blodgett JM, Theou O, Sun MH, Feridooni HA, Mitnitski A, et al. A Frailty Index Based on Deficit Accumulation Quantifies Mortality Risk in Humans and in Mice. Sci Rep. 2017;21.

12. Sampson EL, Blanchard MR, Jones L, Tookman A, King M. Dementia in the acute hospital: prospective cohort study of prevalence and mortality. Br J Psychiatry. 2009;195:61–66.

13. Folstein MF, Folstein SE, McHugh PR. “Mini-mental state”. A practical method for grading the cognitive state of patients for the clinician. Journal of psychiatric research. 1975;12:189–198.

14. Inouye SK, van Dyck CH, Alessi CA, Balkin S, Siegal AP, Horwitz RI. Clarifying confusion: the confusion assessment method. A new method for detection of delirium. Annals of Internal Medicine. 1990;113:941–948.

15. Searle SD, Mitnitski A, Gahbauer EA, Gill TM, Rockwood K. A standard procedure for creating a frailty index. BMC Geriatr. 2008;8:24.

16. Durand CP. Does Raising Type 1 Error Rate Improve Power to Detect Interactions in Linear Regression Models? A Simulation Study. PLOS ONE. 2013;8:e71079.

17. Canevelli M, Cesari M, Francesco R, Trebbastoni, A., Quarata F, Vico C, et al. Promoting the Assessment of Frailty in the Clinical Approach to Cognitive Disorders. Front Aging NeuroSci. 2017;9:36.

18. Blodgett JM, Theou O, Howlett SE, Wu FC, Rockwood K. A frailty index based on laboratory deficits in community-dwelling men predicted their risk of adverse health outcomes. Age Ageing. 2016;45:463–468.

19. Drubbel I, de Wit NJ, Bleijenberg N, Eijkemans RJ, Schuurmans MJ, Numans ME. Prediction of adverse health outcomes in older people using a frailty index based on routine primary care data. J Gerontol A Biol Sci Med Sci. 2013;68:301–308.

20. Pitkala KH, Laurila JV, Strandberg TE, Tilvis RS. Prognostic Significance of Delirium in Frail Older People. Dementia and Geriatrics Cognitive Disorders. 2005;19:158–163.

21. Eeles EM, White SV, O'Mahony SM, Bayer AJ, Hubbard RE. The impact of frailty and delirium on mortality in older inpatients. Age Ageing. 2012;41:412–416.

22. Fong TG, Davis D, Growdon ME, Albuquerque A, Inouye SK. The interface between delirium and dementia in elderly adults. The Lancet Neurology. 2015;14:823–832.

